# The role of reactive oxygen species and calcium signaling in antiviral defense in Arabidopsis

**DOI:** 10.64898/2026.03.03.709233

**Authors:** Jahed Ahmed, Brian Vue, Emily Tipper, Marie Morlans, Nuno Leitão, Nicola Cook, Nathalie Arvy, Arthur Poitout, Marie-Dominique Jolivet, Terezinha Robbe, Stéphanie Pateyron, Christine Payant-Le-Roux, Marie Boudsocq, Alexandre Martinière, Sylvie German-Retana, Myriam Charpentier, Sébastien Mongrand, Véronique Germain, Emeline Teyssier

## Abstract

Plant viruses interfere with host signaling pathways, but it remains unclear how calcium (Ca^2+^) signaling, reactive oxygen species (ROS), and changes in the plasma membrane interact during viral infection. Here, we investigated how plantago asiatica mosaic virus (PlAMV) modulates host Ca^2+^ and ROS-associated signaling in *Arabidopsis thaliana*. Using live-cell imaging and the R-GECO1.2 Ca^2+^ sensor, we observed a rapid increase in cytoplasmic Ca^2+^ before the virus was detected, indicating that Ca^2+^ release occurs early in infection. Genetic analysis showed that GLR, CPK3, and CNGC, core components of Ca^2+^ signaling, limit PlAMV spread between cells, while the usual pattern-triggered immunity (PTI) co-receptors were not needed. This means that Ca^2+^-based antiviral restriction operates independently of PTI. With the plasma membrane-tethered and cytosolic HyPer7 biosensor, we found that ROS levels were lower inside infection foci in the inoculated leaves, but higher in nearby cells, respectively. The NADPH oxidases RBOHD and RBOHF, which produce ROS, slowed down the local viral propagation. The PM sphingolipid biosynthetic enzyme MOCA1 altered ROS patterns and reduced the virus’s spread. Epistasis analysis revealed a functional interaction between RBOHD and MOCA1, suggesting that ROS signaling and plasma membrane sphingolipid homeostasis are interconnected in antiviral defense. Overall, our findings suggest that PlAMV triggers Ca^2+^ influx and ROS signaling at the plasma membrane, which induces sphingolipid reorganization and helps restrict the propagation of the virus. This study shows how Ca^2+^, ROS, and membrane sphingolipid signaling work together in plant antiviral immunity and points to possible ways to improve resistance to viruses.

## Introduction

Plant viruses are obligate intracellular pathogens that pose a significant threat to global agriculture, causing devastating losses in economically important crops and representing a major source of emerging plant diseases (Sanfaçon, 2017). To ensure food security, understanding the molecular mechanisms underlying plant antiviral defenses is crucial for developing virus-resistant crops. Upon infection, viruses hijack host cellular machinery to replicate and spread, moving cell-to-cell through plasmodesmata (PD), plasma membrane (PM)-lined channels modified by viral movement proteins before entering the vasculature for systemic infection (Reagan & Burch-Smith, 2020). Plants mainly protect themselves against viruses through RNA interference (RNAi), which targets viral double-stranded RNA, and Effector-Triggered Immunity (ETI), in which resistance (R) proteins detect viral effectors and trigger defense responses. Recent studies show that RNA viruses, such as plum pox virus, can also trigger Pathogen-Associated Molecular Pattern (PAMP)-Triggered Immunity (PTI). This extra defense begins when Pattern Recognition Receptors (PRRs) detect PAMPs outside the cell (Kørner *et al*., 2013; Nicaise & Candresse, 2017; Teixeira *et al*., 2019).

PTI is well understood in its response to bacterial and fungal infections, but its role in defending against viruses is described (Meier *et al*., 2019, Nicaise *et al*., 2017, Macho and Lozano-Duran 2019). PTI depends on signaling events with reactive oxygen species (ROS) and Ca²⁺ fluxes. These act as important second messengers that control later defense responses, such as PD closure (Cheval & Faulkner, 2018; Wu *et al*., 2020; Köster *et al*., 2022; Tee *et al*., 2022). The PD dynamics regulation by ROS and Ca²⁺ depends on the PD-localized proteins PDLP1 and PDLP5 (Fichman *et al*., 2021; Tee *et al*., 2022). Although ROS and Ca²⁺ signaling during viral infection remains poorly characterized, emerging evidence suggests their critical involvement. For instance, Respiratory Burst Oxidase Homologs (RBOHs), which generate ROS at the PM, influence viral susceptibility or resistance to *red clover necrotic mosaic virus* (RCNMV) or turnip mosaic virus (TuMV), respectively (Hyodo *et al*., 2017; Otulak-Kozieł *et al*., 2020). RCNMV exploits the host ROS machinery by manipulating the immune-related protein RACK1 (Hyodo *et al*., 2017; Alazem & Burch-Smith, 2024). Further supporting the role of Ca²⁺ in antiviral responses, we observed enhanced viral spread in Arabidopsis mutants deficient in the Ca^2+^-Dependent Protein Kinase 3 (CPK3). Strikingly, viral infection alters CPK3 membrane mobility and nanoscale organization within the PM (Jolivet *et al*., 2025), underscoring the importance of PM dynamics in antiviral immunity.

It remains a major challenge in plant immunity research to understand how plant cells integrate Ca²⁺ and ROS signaling during viral infection. While earlier studies have shown that both pathways play a role in antiviral defense, it has been difficult to directly observe Ca²⁺ and ROS activity in living plant cells during viral attack. In this study, we used live-cell biosensors for the first time to track Ca²⁺ and ROS signals in response to plant virus infection. Using the genetically encoded fluorescent Ca²⁺ sensor R-GECO1.2 and both cytosolic and plasma membrane-tethered ROS biosensors HyPer7, we showed spatiotemporal patterns of signaling events triggered by *plantago asiatica mosaic virus* (PlAMV) infection in *Arabidopsis thaliana*.

PlAMV, a potexvirus causing lily necrosis serves as a model virus (Hashimoto *et al*., 2016; Komatsu & Hammond, 2022). Potexviruses, including agronomically significant pathogens such as pepino mosaic virus (PepMV), are positive strand RNA viruses whose genome encodes three movement proteins (TGBp1-3), a coat protein (CP), and an RNA-dependent RNA polymerase (RdRp). Our previous work with fluorescently tagged PlAMV enabled quantitative tracking of viral spread via epifluorescence microscopy and whole-plant imaging (Jolivet *et al*., 2025) and revealed CPK3 and REMORIN reorganization during infection (Jolivet *et al*., 2025). Here we use Arabidopsis/PlAMV pathosystem to show that early Ca²⁺ influx and changes in ROS levels act together to limit viral infection.

## Methods and materials

### Plant material and genetic crosses

The Arabidopsis mutant in Col-0 background were previously described: *cpk3* (Mehlmer *et al*., 2010), *glr3.2/3.6* (Toyota *et al*., 2018), *moca1* (Jiang *et al*., 2019), *cngc19* (Meena *et al*., 2019), *osca1;1/1;2/1;3* (kindly provided by Dr. Cyril Zipfel, personal communication), *rbohC* (Foreman *et al*., 2003), *rbohD* (Torres *et al*., 2002), *rbohF* (Torres *et al*., 2002), *dgk5* (Kalachova *et al*., 2022), *hpca1* (Wu *et al*., 2020), *serk1* (Heese *et al*., 2007), *bik1* (Veronese *et al*., 2006), *bik1;5/bkk1-1* (He *et al*., 2007). To monitor intracellular reactive oxygen species (ROS) accumulation, *Arabidopsis thaliana* lines expressing the genetically encoded biosensor HyPer7 (Ugalde *et al*., 2021) were used. The HyPer7 sensor was backcrossed into mutant backgrounds (*moca1*, *rbohD and cpk3-2*) through genetic crosses. Briefly, homozygous mutant lines were crossed with the HyPer7-expressing line, and F2 progeny were screened for homozygous mutants carrying the sensor. To monitor Ca^2+^ dynamic, *Arabidopsis thaliana* (ecotype Col-0) plants were used for stable transformation. The binary vector pNOS:BAR:TNOS – LjUBI1:RGECO1.2:T35S was generated via GoldenGate cloning (Weber *et al*., 2011) and consist of the phosphinothricin acetyltransferase (BAR) driven by the nopaline synthase promoter (pNOS) and terminated by the NOS terminator (TNOS) for plant selection. The Ca^2+^ indicator R-GECO1.2 was cloned under the control of the *Lotus japonicus* ubiquitin promoter (LjUBI1) and terminated by the cauliflower mosaic virus 35S terminator (T35S). The binary vector pNOS:BAR:TNOS – LjUBI1:RGECO1.2:T35S was introduced into the *Agrobacterium tumefaciens* GV3101 (Lieasible, NY, USA), and stable transgenic *A. thaliana* expressing R-GECO1.2 were generated using the floral dip method (Zhang *et al*., 2006). T1 plants were selected on Murashige and Skoog medium supplemented with 10 mg L^-1^ glufosinate ammonium for selection of BAR-resistant transformants. Resistant T1 seedlings were screened for expression of R-GECO1.2 by fluorescence microscopy. Homozygous T4 lines were used for subsequent experiments.

### RNA-seq analysis

Three-week-old *A. thaliana* leaves were infiltrated with PlAMV-GFP or free-GFP as a control. Two leaves of two individual plants were harvested per condition and per genotype. Three biological repeats were performed.

Total RNAs were extracted using the Qiagen RNeasy Plant Mini Kit, followed by DNAse treatment, according to manufacturer’s instructions and were further purified using the RNA Clean & Concentrator-5 Kits (Zymo Research). RNA integrity was checked on Agilent RNA Nano chip and quantified using Quant-iT™ RiboGreen (Thermo Fisher Scientific Inc).

RNA-seq libraries were constructed with 250ng of total RNA using the QuantSeq 3’ mRNA-Seq Library Prep Kit (FWD) for Illumina 96 preps (Clinisciences) according to the supplier’s instructions. Libraries quality were checked on Agilent DNA HS chip and quantified with Quant-iT™ PicoGreen™ dsDNA Reagent (Thermo Fisher Scientific Inc) to generate an equimolar pool. Libraries were sequenced in single-end (SE) mode with 150 bases for each read on our NextSeq500 (Illumina).

Read 2s were excluded from the pre-processing. UMIs were removed and appended to the read identifier with the extract command of UMI-tools (v1.0.1, Smith *et al*., 2017). Reads where any UMI base quality score falls below 10 were removed. To remove adapter sequences, poly(A), poly(G) sequences, and low quality low-quality nucleotides, reads were trimmed with BBduk from the BBmap suite (v38.84, Bushnell and Brian, 2014) with the options k=13 ktrim=r useshortkmers=t mink=5 qtrim=r trimq=10 minlength=30. Trimmed reads were then mapped using STAR (v2.7.3a, Dobin et al. 2013), with the following parameters --alignIntronMin 5 --alignIntronMax 60000 --alignMatesGapMax 6000 –alignEndsType Local--outFilterMultimapNmax 20 --outFilterMultimapScoreRange 0 –outSAMprimaryFlag AllBestScore –mismatchNoverLmax 0,6 on the *Arabidopsis thaliana* genome reference (TAIR10 www.arabidopsis.org). Reads with identical mapping coordinates and UMI sequences were collapsed to remove PCR duplicates using the dedup command of UMI-tools with the default directional method parameter. RSeQC (v2.6.6, Wang *et al*., 2012) was used to evaluate deduplicated mapped reads distribution. Deduplicated reads were counted using HTSeq-count (v0.12.4, Anders *et al*., 2015) (htseq-count mode intersection-nonempty) based on the gene annotations. Between 10,5 (84,1% of raw reads) and 12,1 (87,7% of raw reads) millions of deduplicated PE/SE reads were associated to annotated genes.

Statistical analyses were conducted on R 4.3.3 (R Core Team, 2020) using the R script-based tool DiCoExpress (Lambert *et al*., 2020; Baudry *et al*., 2022) based on the Bioconductor package edgeR (v 4.0.16, (Robinson *et al*., 2010; McCarthy *et al*., 2012). Genes with low counts were filtered using the “filterByExpr” function where the group argument specifies the biological conditions (free-GFP or PlAMV-GFP), the min.count value is set to 3 and the other arguments are set to their default values. Libraries were normalized with the Trimmed Mean of M-values (TMM) method with the default parameter values.

The differential analysis was based on a negative binomial generalized linear model in which the logarithm of the average gene expression is an additive function of a condition effect (free-GFP or PlAMV-GFP), and a replicate effect (3 modalities) and we tested the difference between two conditions using a likelihood ratio test. Raw p-values were adjusted with the Benjamini–Hochberg procedure to control the false discovery rate. We checked that the distribution of the raw p-values followed the quality criterion described by Rigaill *et al., 2018* and declared that a gene was declared differentially expressed if its adjusted p-value was lower than 0.01.

GO enrichment was analyzed with Panther using Panther GO-slim biological processes and cellular component.

### Local viral propagation assay

The assay was conducted as previously described (Jolivet *et al*., 2025) using either PlAMV-GFP, or PlAMV-mCherry, respectively GFP- and mCherry-labeled infectious clones of PlAMV that can be delivered via agroinfiltration (Minato et al., 2014). *Agrobacterium tumefaciens* strain GV3101, transformed with PlAMV-GFP and/or PlAMV-mCherry, was infiltrated into 3-week-old *Arabidopsis thaliana* plants at an optical density (OD600) of 0.150 for PlAMV-GFP and 0.100 for PlAMV-mCherry. Viral spread was monitored 5 days post-infection using an Axiozoom V16 macroscope to quantify the surface of viral GFP-labelled infection foci on the inoculated leaves. Infection foci surface was quantified automatically with a custom macro in Fiji software (http://fiji.sc/). Statistical analysis was performed using two-way ANOVA with Dunnett’s and Tukey’s post-hoc test for multiple comparisons.

### Confocal microscopy and image acquisition

Leaves infected with PLAMV-mCherry were excised from 3-week-old *A. thaliana* plants expressing the biosensor HyPer7-KRas. The leaves were then rested for 10 minutes to avoid wounding-induced ROS production. Subsequently, intracellular ROS was visualized in the infected areas and neighboring cells. Intracellular ROS accumulation was visualized using a ZEISS LSM 880 confocal laser scanning microscope with a 20× objective. The HyPer7 sensor was excited at 488 nm (oxidized form) and 405 nm (reduced form), with emission collected between 508–535 nm. The 488/405 nm fluorescence ratio was calculated to determine ROS levels, with acquisition parameters kept consistent across experiments. For each experiment, 10–12 regions of interest (ROIs) per image were randomly selected for quantification. The fluorescence ratio (488/405 nm) was calculated for HyPer7 signals. Signal quantification and ratiometric image analysis were performed using FIJI (Schindelin *et al*., 2012). Data were collected from at least three independent biological replicates, with three leaves per replicate (each from a separate plants). Statistical analysis was performed using GraphPad Prism 9 (GraphPad Software, San Diego, CA, USA). Data were analyzed by one-way ANOVA followed by Dunnett’s multiple comparisons test. Results are presented as mean ± SEM (standard error of the mean) of at least three independent biological replicates. Differences were considered statistically significant with at least *p < 0.05.

### Ca^2+^ imaging in R-GECO Arabidopsis leaves

Three-week-old *Arabidopsis thaliana* (ecotype Col-0) plants expressing R-GECO1.2 were grown under controlled environment conditions (22 °C day, 20 °C night, 16-hour photoperiod, 80% relative humidity). Leaves were infiltrated with *Agrobacterium tumefaciens* GV3101 carrying either free-GFP (“mock” control) or PlAMV-GFP constructs. *Agrobacterium* cultures were grown overnight in liquid LB medium containing kanamycin (50 µg ml^-1^) gentamicin (25 µg ml^-1^) and rifampicin (50 µg ml^-1^) at 28°C with shaking (200 rpm). Bacterial culture were centrifuged and resuspended in infiltration buffer (10 mM MgCl₂, 10 mM MES pH 5.6, 100 µM acetosyringone) to a final OD₆₀₀ of 0.1. Leaves were imaged at 3 days post infiltration using a Zeiss LSM880 confocal laser scanning microscope. Z-stacks comprising three optical sections were acquired using sequential excitation to avoid spectral overlap between GFP and R-GECO1.2. GFP was excited at 488 nm and detected between 500–550 nm, while R-GECO1.2 was excited at 561 nm and detected between 580–650 nm. Time-series images were collected at 5-min intervals over a 3-h period. Image processing and analysis were performed using Fiji/ImageJ. Normalized datasets (ΔF/F) were calculated as (F - F_0_)/F_0_, where F_0_ represents the average of at least 10 min of baseline values.

## Results

### PTI-related genes are not required for restriction of PlAMV propagation

We first determine whether PlAMV propagation is altered in Arabidopsis mutants defective in key PTI-related genes. The somatic embryogenesis receptor-like kinases (SERKs), including BAK1 (BRI1-ASSOCIATED KINASE1), BKK1 (BAK1-LIKE 1), and SERK1, are well-characterized plasma membrane co-receptors that associate with pattern-recognition receptors (PRRs) to initiate PTI. Downstream of these complexes, the receptor-like cytoplasmic kinase BIK1 acts as a key signaling mediator linking PRR activation to early defense outputs such as Ca^2+^ influx, ROS production, and MAPK activation (Jiang *et al*., 2019a). Analyses of PlAMV propagation in single *serk1*, *bik1* and in the double mutant *bak1;5/bkk1-1* revealed no significant difference in viral spread compared to wild-type Col-0 plants (Figure S1), indicating that these co-receptors are not directly involved in PlAMV propagation.

### Transcriptomic analysis upon PlAMV infection

To study how PlAMV affects host signaling, we compared the transcriptomes of infected leaf areas containing the virus with similar regions from healthy leaves. Considering a threshold p-value < 0.01, the RNAseq analysis revealed 739 up-regulated genes and 562 down-regulated genes by PlAMV infection (Figure S2A, Table S1).

Gene Ontology (GO) analysis of up-regulated genes highlighted significant enrichment of biological processes associated with oxidative signaling and stress responses (Fig. S2B). Among the top categories, “response to reactive oxygen species” was highly overrepresented, consistent with elevated oxidative stress signaling following PlAMV infection. GO enrichment of biological processes associated with the down-regulated genes highlighted defense and stress responses, signal transduction, as well as Ca^2+^-mediated signaling and regulation of cell communication, suggesting that the virus may manipulate plant signaling to bypass defense and spread (Figure S2C). Genes involved in some basic cellular functions are also repressed (Fig. S2C),, suggesting that the plant shifts from growth to defense during infection. GO analysis of cellular components for downregulated genes reveals that membrane-related compartments are particularly affected (Fig. S2D), indicating that PlAMV infection strongly affects membrane-related cellular structures. We further looked at the genes involved in Ca^2+^ signaling (Figure S2E). A subset of genes was significantly upregulated following PlAMV infection, including several Ca^2+^ sensors protein kinases from CIPK and CPK families as well as Ca^2+^ channels and transporters from OSCA, CNGC and GLR families. These genes are known to participate in Ca²⁺ influx, homeostasis, and signaling during stress and pathogen responses (Ming *et al*., 2025), suggesting enhanced activation of specific Ca²⁺ signaling branches in infected tissues. Conversely, a larger group of Ca²⁺-associated genes were downregulated in response to PlAMV infection. These included several Ca^2+^ sensors from CIPK and CPK families, as well as from CAM (calmodulin) and CML (calmodulin-like) sensor relay families. PlAMV also repressed the expression of Ca^2+^ channels and transporters including *CNGC19*. Interestingly, *RBOHD* was also downregulated in infection foci. Collectively, these results suggest that PlAMV infection triggers a transcriptional reprogramming of host genes associated with ROS generation, H₂O₂ response, and Ca^2+^ signaling, reflecting a coordinated activation of redox-dependent defense and signaling pathways during PlAMV infection.

### Monitoring Ca^2+^ during virus propagation

To investigate the role of cytoplasmic Ca²⁺ signaling in PlAMV movement, we monitored Ca²⁺ dynamics and viral spread in *Arabidopsis thaliana* epidermal cells using a dual-imaging approach. Transgenic plants expressing the red fluorescent Ca²⁺ sensor R-GECO1.2 were infiltrated with PlAMV-GFP, and fluorescence signals were recorded over time inside the infection foci (local area) and outside the infection foci at 3-4 cells of distance (distal area, not-yet-infected cells). We readily observed that non-infected cells in the distal area exhibited a marked increase in R-GECO1.2 fluorescence intensity over time, indicating a rapid rise in cytoplasmic Ca²⁺ levels (Figure 1, Figures S3, S4 for replicates and movies 1, 2, 3). Notably, the Ca²⁺ signal was first detected at the infection site and subsequently propagated to not-yet-infected neighboring cells. Importantly, the elevation in cytosolic Ca²⁺ preceded the appearance of the PlAMV-GFP signal in the distal area, which became detectable only after an approximate 30 min delay (Figure 1C, S3A, S4A and movies 1, 2 and 3). Quantitative analysis of fluorescence intensities within a defined region of interest (ROI) confirmed that Ca²⁺ signaling occurred prior to viral accumulation (Figure 1B, 1C). The R-GECO1.2 signal peaked earlier in the time course, whereas the PlAMV-GFP signal showed a delayed but steady increase, consistent with viral propagation. Notably, only one wave of Ca²⁺ is observed during more than 3 hours of imaging. As control, this response was not observed in cells expressing free GFP, indicating that the Ca²⁺ elevation is induced by PlAMV infection (Figure S5). These findings suggest that cytoplasmic Ca²⁺ release is an early event during PlAMV infection and suggest that Ca²⁺ signaling may facilitate early plant stress response or prime host cells for subsequent viral propagation.

**Figure 1.**
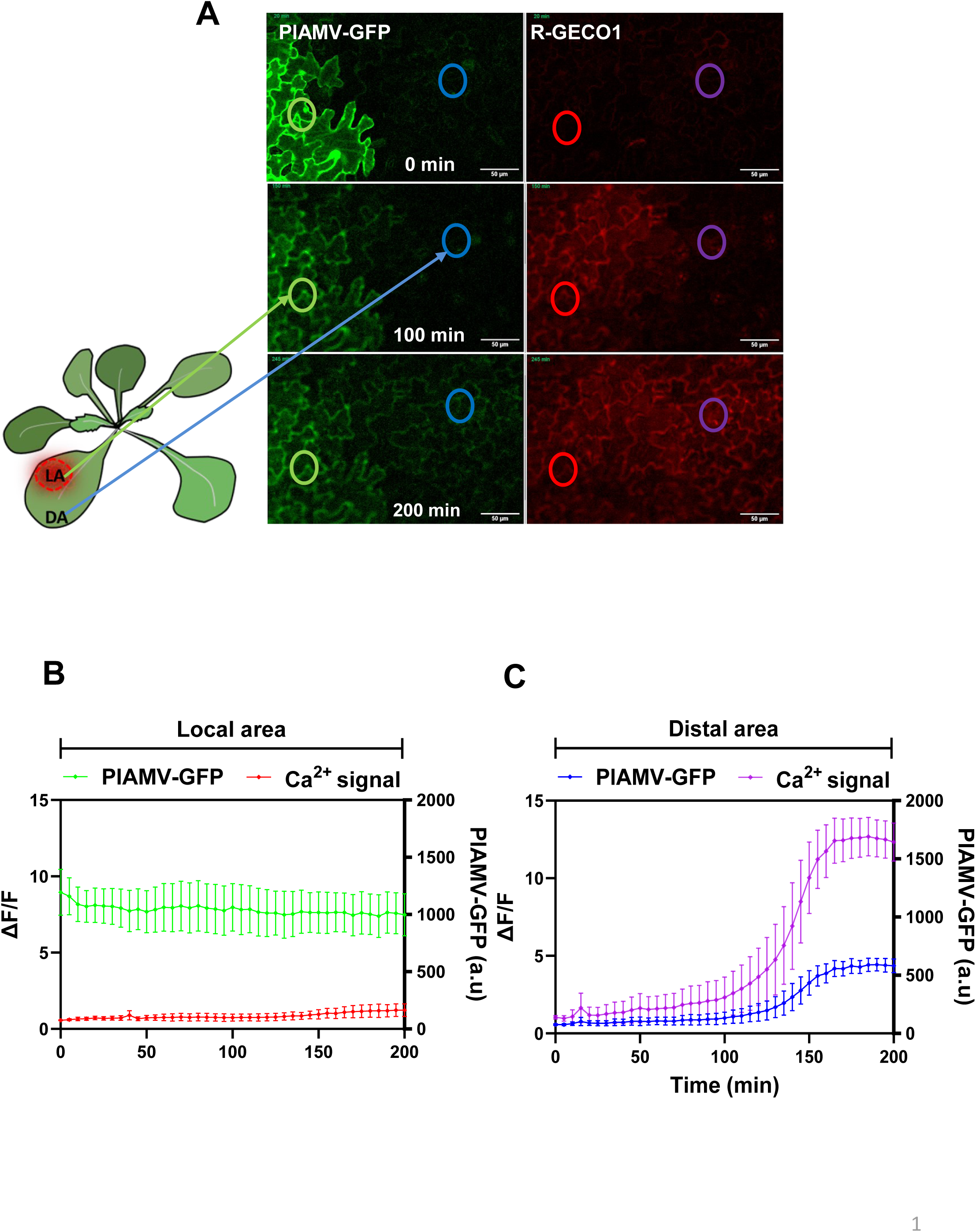
Cytoplasmic Ca^2+^ release paves the way of PlAMV propagation. **A**, Representative confocal images of *A. thaliana* pavement cells stably expressing the red genetically encoded Ca^2+^ indicator version 1.2 (R-GECO1.2) and agro-infiltrated with PlAMV-GFP. Images in the left panels show the recording of the PlAMV-GFP signal, whereas the right panels display the simultaneous acquisition of red fluorescence channel corresponding to R-GECO1.2 signals. The acquisition is performed from a cell already infected (local area, LA), from which the calcium signal propagates to neighboring cells (distal area, DA) prior to the movement of the PlAMV-GFP. Time stamps are indicated in minutes. The region of interest (ROI) is marked by a circle. Scale bars represent 50 µm. **B**, Quantification of fluorescence intensity over time of R-GECO1.2 (red) and PlAMV-GFP (green) within the ROI corresponding to the LA (A). **C**, Quantification of fluorescence intensity over time of R-GECO1.2 (violet) and PlAMV-GFP (blue) within the ROI corresponding to the DA (A).

### GLR, CPK3, and CNGC are involved in restricting PlAMV propagation in *Arabidopsis thaliana*

To assess the involvement of Ca²⁺-associated signaling components in PlAMV infection, we performed the viral propagation assay in *Arabidopsis thaliana* mutants defective in several Ca^2+^ channels. *GLR3.3* and *GLR3.6* encode glutamate receptor-like channels implicated in Ca²⁺-mediated signaling and long-distance electrical responses (Mousavi *et al*., 2013 and Toyota *et al*., 2018). *CNGC19* encodes a cyclic nucleotide-gated ion channel contributing to Ca²⁺ influx during defense activation (Meena *et al*., 2019). OSCA1s are reduced hyperosmolality-induced [Ca^2+^]i increase channels that contribute to Ca²⁺ influx in response to osmotic stresses and in stomatal immunity (Yuan *et al*., 2014, Thor *et al*., 2020). We also analyzed the mutant of *CPK3* as a positive control, which encodes a calcium-dependent protein kinase involved in stress and immune signal transduction including viral propagation (Jolivet *et al*., 2025). Leaves were agroinoculated with PlAMV-GFP, and infection foci were visualized under fluorescence microscopy at 5 days post-inoculation (dpi). Successful viral infection and spread were marked by GFP fluorescence. As shown in Figure 2A, PlAMV-GFP infection foci appeared markedly larger in the *glr3.3/3.6, cngc19* and *cpk3* mutant backgrounds compared to the wild-type (Col-0), indicating enhanced viral propagation in these lines. By contrast, *osca1;1/1;2/1;3* exhibited wild-type area, suggesting that those OSCA1 channels are not involved. To quantify these observations, the area of GFP-positive infection foci was measured and normalized to the mean foci area in Col-0 plants (Figure 2B). Statistical analysis confirmed that the *glr3.3/3.6*, *cngc19* and *cpk3* mutants exhibited significantly larger infection foci relative to Col-0, while *osca1;1/1;2/1;3* displayed similar infection foci. These findings indicate that GLRs, CPK3 and CNGC19, but not OSCA1s, play a significant role in limiting PlAMV cell-to-cell propagation in *A. thaliana*. Together, these results demonstrate the importance of a subset of calcium-signaling components in the spatial restriction of PlAMV during early stages of infection.

**Figure 2.**
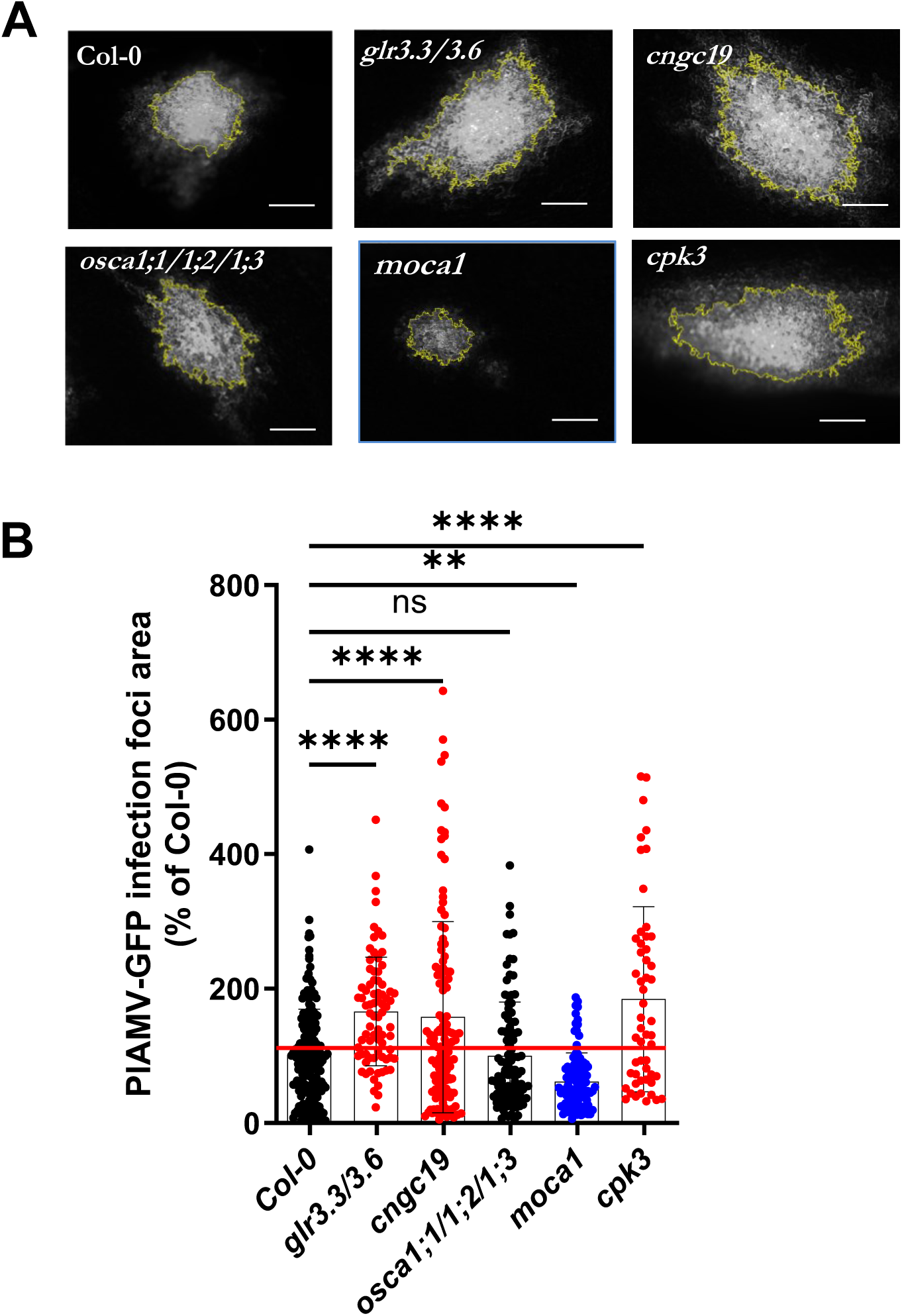
*Arabidopsis thaliana* calcium-related genes (*GLR, CPK3, MOCA1* and *CNGC*) are specifically involved in the restriction of PlAMV propagation but not OSCA1s. A, Representative images of PlAMV-GFP infection foci at 5 dpi in the different calcium-related mutant backgrounds. Scale bar = 500 µm. **B**, Graph represents the mean area of PlAMV-GFP infection foci 5 dpi in different calcium-related mutant lines, normalized and expressed as a percentage of PlAMV-GFP propagation in Col-0. Three independent biological repeats were performed, with at least 50 infection foci per experiment and per genotype. Significant differences were revealed using a one-way ANOVA followed by a Dunnett’s multiple comparison test (**** p <0.001, ** p <). ns: not significant.

### PlAMV local propagation is hampered in Arabidopsis mutant *monocation-induced [Ca^2+^]_i_ increases 1 (moca1)*

To further investigate the role of Ca^2+^ signaling in viral plant immunity, we analyzed the *A. thaliana* mutant *monocation-induced [Ca^2+^*]i *increases 1* (*moca1*). This mutant was originally isolated through Ca^2+^-imaging-based forward genetic screens conducted under salt stress (Jiang *et al*., 2019b). MOCA1 was further characterized as a hypormorphic mutant of the glucuronosyltransferase IPUT1 (Inositol phosphorylceramide glucuronosyltransferase 1). IPUT1 catalyzes the first glycosylation step in the biosynthesis of glycosyl inositol phosphorylceramides (GIPCs), which constitute the predominant class of sphingolipids in the plant plasma membrane. IPUT1 was shown to be required for salt-induced depolarization of the cell-surface potential, Ca^2+^ spikes and waves, and Na^+^/H^+^ antiporter activation (Jiang *et al*., 2019b). Na^+^ binds to GIPC negatively charged polar head to gate Ca^2+^ influx channels. This salt-sensing mechanism might imply that PM sphingolipids are involved in adaptation to various environmental stress. Thus, we assessed PlAMV-GFP virus spread at 5 days dpi in *moca1*. Strikingly, *moca1* mutant exhibited a strongly reduced PlAMV-GFP infection foci compared to wild-type Col-0 as shown in Figure 2, suggesting a role of IPUT1 as a susceptibility factor of viral propagation.

### PM-bound ROS-producing NADPH oxidases RBOHD and RBOHF restrict PlAMV propagation

To further assess the role of ROS production during viral infection, we examined *A. thaliana* mutants impaired in ROS-generating NADPH oxidases in leaves, namely Respiratory Burst Oxidase Homolog RBOHD and RBOHF (Remans *et al*., 2010; Morales *et al*., 2016), using a PlAMV-GFP. At 5 dpi, plants carrying mutations in RBOHD or RBOHF displayed significantly larger infection foci compared to Col-0 (Figure 3), suggesting compromised viral containment. By contrast, plants mutated in *RBOHC*, which is not expressed in leaves (Remans *et al*., 2010; Morales *et al*., 2016), displayed wild-type response. DGK5 (diacylglycerol kinase 5) has been shown to activate RBOHD and positively regulate pattern-triggered immunity (PTI) (Kong *et al*., 2024), while HPCA1 (hydrogen-peroxide-induced Ca^2+^ increases 1) encodes a leucine-rich-repeat receptor kinase notably involved in systemic cell-to-cell ROS signaling in response to a local bacterial infection (Fichman *et al*., 2022). We thus examined whether DGK5 and HPCA1 also contribute to antiviral defense. The *dgk5* mutant displayed significantly larger infection foci than Col-0 (Figure 3), indicating that DGK5-dependent activation of RBOHD is critical for restricting PlAMV local propagation. In contrast, infection foci in the *hpca1* mutant were comparable in size to those of Col-0, indicating that HPCA1 does not contribute to this antiviral mechanism. Together, these results identify RBOHD and RBOHF as key ROS-producing enzymes that restrict local PlAMV propagation.

**Figure 3.**
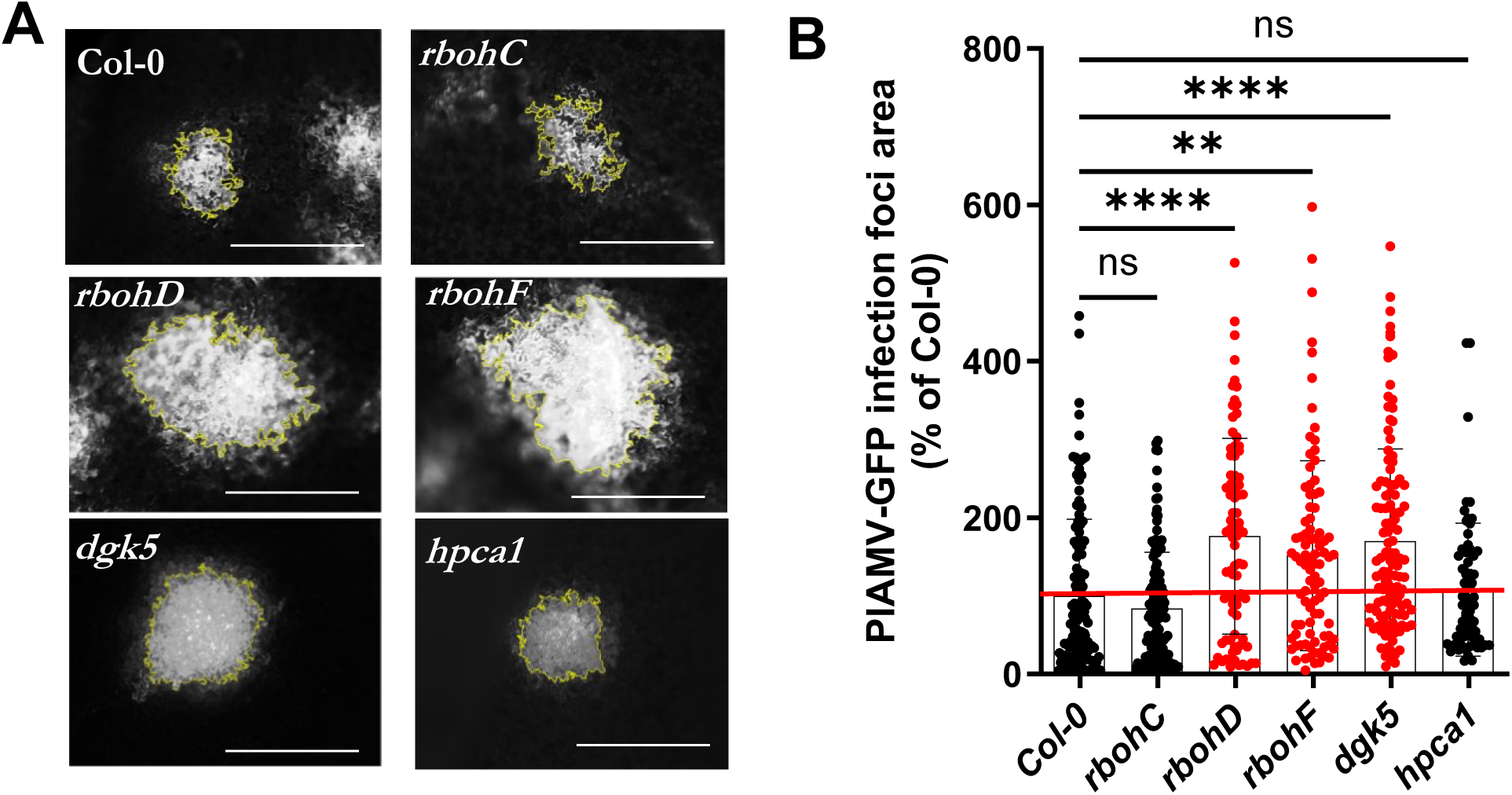
*Arabidopsis thaliana* ROS-related genes (*RBOHD and RBOHF*) are specifically involved in the restriction of PlAMV cell-to-cell movement. A, Representative images of PlAMV-GFP infection foci at 5 dpi in the different ROS-related mutant backgrounds. Scale bar = 500 µm. **B**, Graph represents the mean area of PlAMV-GFP infection foci 5 dpi in different ROS-related mutant lines, normalized and expressed as a percentage of PlAMV-GFP propagation in Col-0. Three independent biological repeats were performed, with at least 50 infection foci per experiment and per genotype. Significant differences were revealed using a one-way ANOVA followed by a Dunnett’s multiple comparison test (****p <0.001, ** p <). ns: not significant.

### Plasma membrane-localized and cytosolic ROS accumulate in neighboring not-yet-infected cells during PlAMV infection

To investigate the spatial distribution of reactive oxygen species (ROS) during PlAMV infection, we employed *A. thaliana* lines stably expressing HyPer7-kRas a genetically encoded hydrogen peroxide (H₂O₂) sensor in the cytosol or targeted to the inner leaflet of the plasma membrane (PM) via a kRas farnesylation (Figure S6). Five-days after infection with PlAMV-mCherry, cytosolic or PM-located ROS levels were significantly dampened within the infection foci (local Area) compared to non-infected control plants (Col-0) (Figure 4). By contrast, ROS levels were also significantly elevated in area distal to the infection foci (Distal Area) compared to non-infected controls (Figure 4). These results indicate that H₂O₂ exhibits a similar spatial redistribution pattern in the cytosol and at the PM, characterized by localized suppression within infection foci and accumulation in surrounding, non-infected cells.

**Figure 4.**
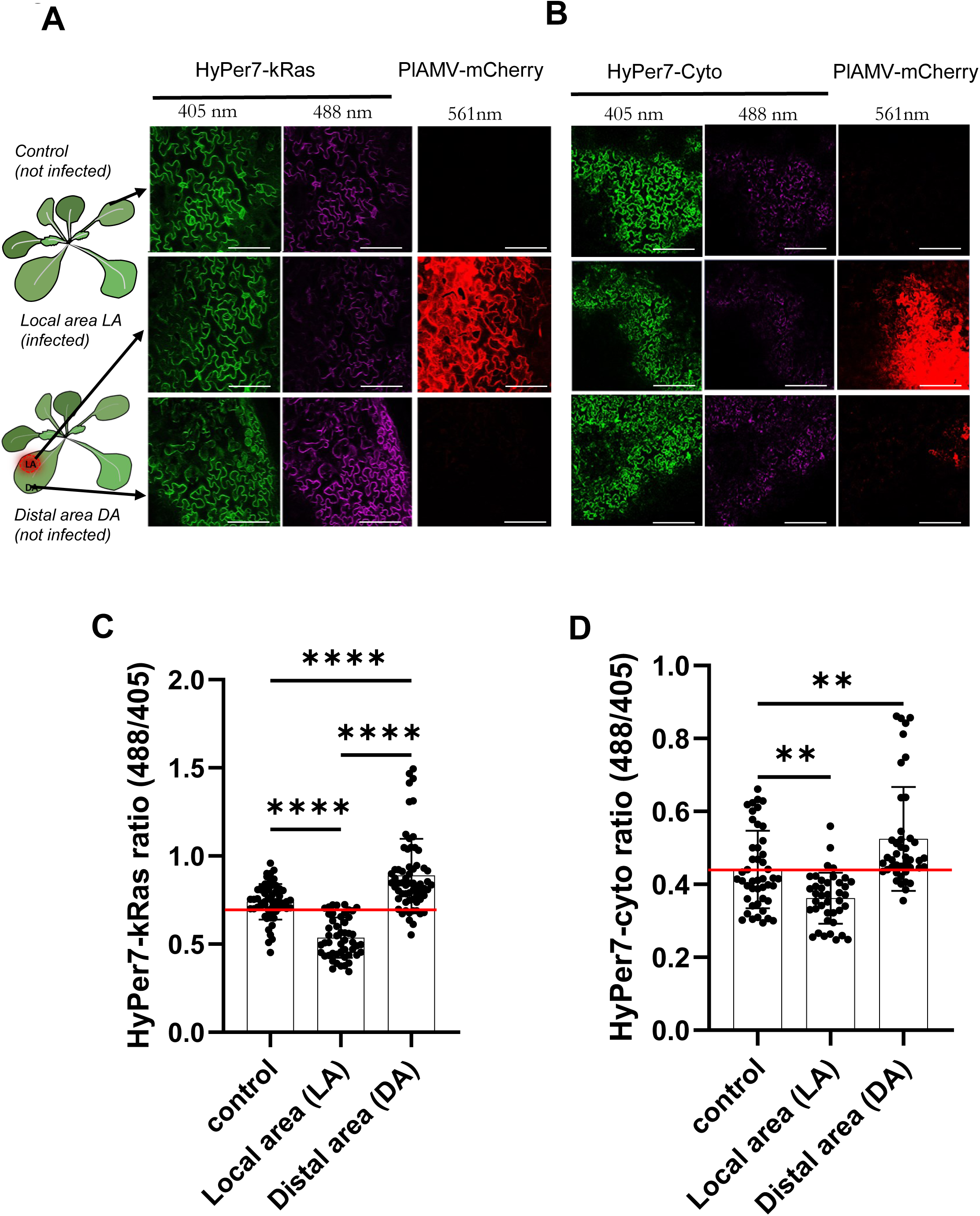
ROS level measured in Arabidopsis expressing plasma membrane-bound HyPer7-kRas and cytosolic HyPer7 infected with PlAMV-mCherry. **A,B**, Schematic representation of inside the foci of infection (Local Area, LA) or neighboring cells (Distal Area, DA), compared to the non-infected control plant for ROS measurements; confocal microscopy images of the reduced form of Hyper7 (green, excited at 405 nm) and oxidized form (magenta, excited at 488 nm), as well as PlAMV-mCherry (detected at 561 nm) at 5 dpi, in stable transgenic line expressing HyPer7-kRas located at the PM (A), and HyPer7-cyto located in the cytosol (B). Scale bar = 100 nm; **C,D**, ROS level was quantified at the PM (C) or in the cytosol (D), inside the foci of infection (Local Area, LA) or neighboring cells (Distal Area, DA), or measured in a non-infected plant (Control). The data were obtained from 3 independent experiments (N=3). Statistical differences were determined by one-way ANOVA followed by Tukey’s multiple comparisons test (*** p <0.001, ** p <).

### Plasma membrane–associated ROS accumulation during PlAMV infection requires MOCA1, RBOHD, and CPK3

We further monitored plasma membrane–associated ROS during PlAMV infection using the HyPer7-kRas biosensor in *A. thaliana* mutants defective in MOCA1, RBOHD, or CPK3. Plants were inoculated with PlAMV-mCherry, and ROS levels were quantified at viral infection foci (local area, LA), surrounding cells (distal area, DA), and in non-infected controls.

In wild-type Col-0 plants, ROS levels rose in the DA relative to uninfected tissue, consistent with viral infection-induced ROS signaling (Figure 4). In contrast, *moca1, rbohD and cpk3-2* mutants failed to modulate ROS either at the LA or DA, showing levels comparable to uninfected controls (Figure S7). This suggests that IPUT1, RBOHD and CPK3 are required to finely tune the PlAMV-induced spatial regulation of ROS levels at the plasma membrane, highlighting their possible roles in coordinating localized oxidative responses during viral infection.

### Study of putative cross talk between ROS and Ca^2+^ signaling during PlAMV infection

To explore a potential cross-talk between RBOHD-mediated ROS production and calcium signaling during PlAMV infection, we generated the double mutants *rbohD/moca1, rbohD/cpk3-2* and *cpk3-2/moca1* and analyzed PlAMV-GFP propagation in the single and double mutants compared to Col-0.

The mean infection foci area were not significantly different between *rbohD/moca1* and *rbohD* (Figure 5A, and B). This result suggests that RBOHD and MOCA1/IPUT1 interfere with PlAMV spread efficiency in the same pathway, RBOHD acting downstream of IPUT1. Indeed, the negative impact of *RBOHD* loss of function overcomes the positive effect of GIPC decrease in *moca1*.

**Figure 5.**
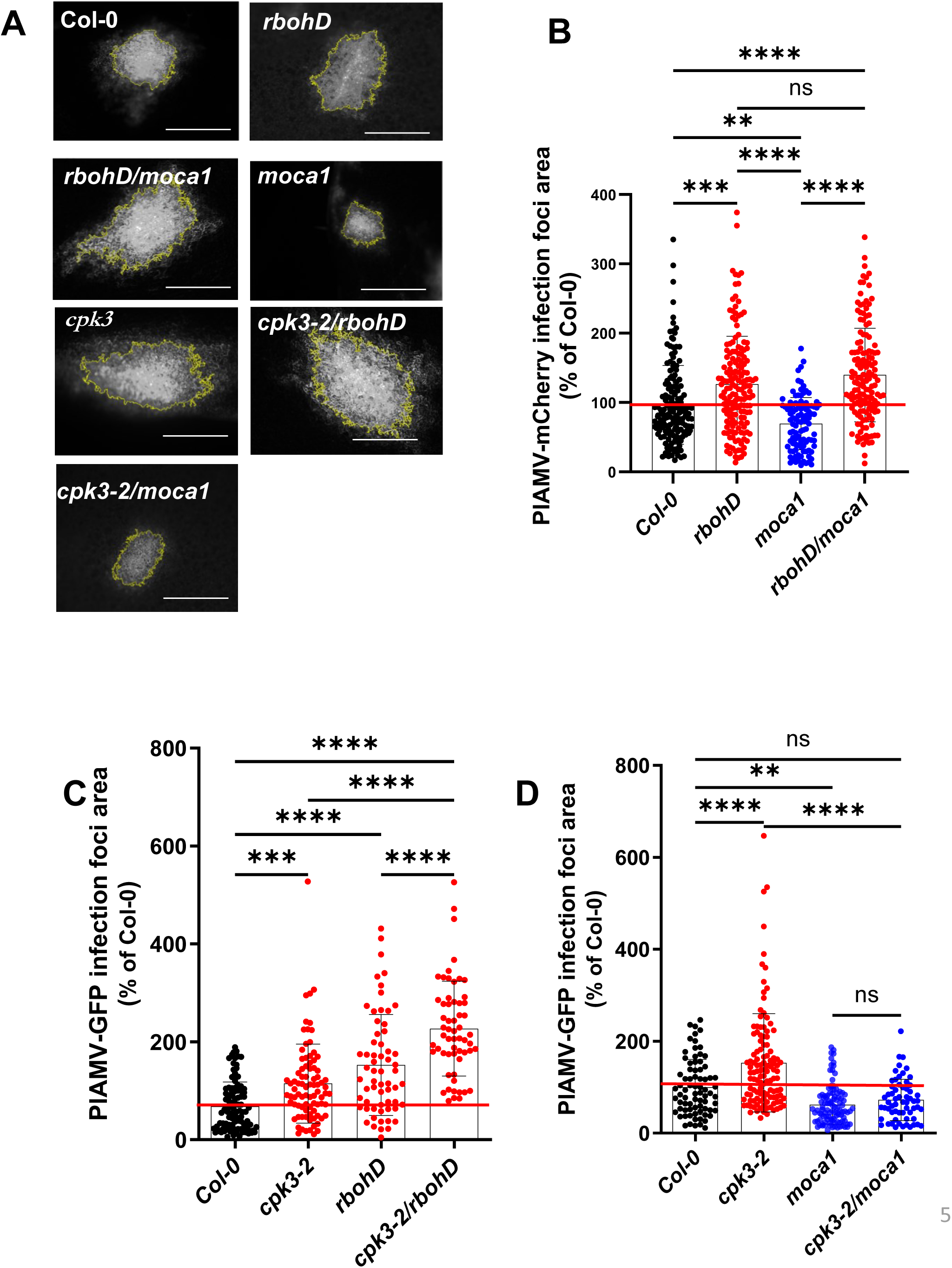
*Arabidopsis thaliana* PM lipid-related gene, *moca1*, *cpk3-2* and *rbohD* crosstalk and their involvement in the restriction of PlAMV propagation. A, Representative images of PlAMV-GFP infection foci at 5 dpi in the different PM lipid-related mutant backgrounds. Scale bar = 500 µm. **B**, Graph represents the mean area of PlAMV-GFP infection foci 5 dpi in PM lipid-related mutant lines and double mutant (*rbohD/moca1*), normalized and expressed as a percentage of PlAMV-GFP propagation in Col-0. Three independent biological repeats were performed, with at least 50 infection foci per experiment and per genotype. Significant differences were revealed using a one-way ANOVA followed by a Tukey’s multiple comparison test (***p <0.001). ns: not significant. **C**, Graph represents the mean relative PlAMV-GFP foci area in wild type (Col-0) and *CPK3*-2 and *RBOHD* mutants from 3 independent experiments (N=3). **D**, PIAMV-GFP infection foci area (expressed as % of Col-0) in wild-type Col-0, *cpk3*-2, *moca1*, and the double mutant *cpk3-2/moca1*. Plants lacking CPK3 showed significantly increased infection foci area compared to Col-0, indicating that CPK3 negatively regulates PIAMV spread. In contrast, *moca1* mutants displayed reduced propagation compared to *cpk3*-2, and the double mutant *cpk3*-2/*moca1* resembled *moca1* rather than *cpk3*-2, demonstrating that MOCA1 is dominant over CPK3 in controlling virus spread. Statistical differences were determined by one-way ANOVA followed by Tukey’s multiple comparisons test (***p <0.001). ns: not significant.

Conversely, virus propagation was significantly increased in the double mutant *rbohD/cpk3-2* compared to single individual *rbohD* and *cpk3-2* mutants (Figure 5A and C), suggesting that RBOHD and CPK3 act in two different additive pathways to mediate antiviral defense.

Finally, PIAMV-GFP propagation in the double mutant *cpk3-2/moca1* was indistinguishable from the *moca1* single mutant indicating that MOCA1/IPUT1 is epistatic to CPK3 in controlling virus spread (Figure 5A and D). Taken together, these results suggest that CPK3 and MOCA1/IPUT1 function together in the regulation of PIAMV spread, with MOCA1/IPUT1 acting downstream of CPK3.

## Discussion

In this study, we examined how a potexvirus interacts with host membrane signaling networks, focusing on the roles of Ca^2+^ dynamics and ROS production, and on how these signaling components work together to control viral spread in *Arabidopsis thaliana*. Our findings suggest that PlAMV may affect host pathways to help infection, while the plant uses specific signaling at the plasma membrane to slow down viral movement.

Mutants defective in canonical pattern-triggered immunity (PTI) components, including SERK co-receptors, did not exhibit altered PlAMV propagation (Figure S1). These data indicate that PlAMV evades canonical PTI pathways. Nonetheless, our transcriptomic analysis shows that PlAMV infection alters Ca²⁺ and reactive oxygen species (ROS) signaling pathways in *Arabidopsis thaliana*. These changes point to a coordinated adjustment of these connected networks. The heatmap of differentially expressed genes shows that some CIPK, OSCA, and GLR family members are upregulated, consistent with activation of Ca²⁺ influx and sensing at the plasma membrane (Yuan *et al*., 2014; Toyota *et al*., 2018). Meanwhile, several CPK, CML, and CNGC genes are downregulated, suggesting that downstream Ca²⁺-dependent signaling components involved in immune responses are selectively changed (Ranty *et al*., 2016; Bredow and Monaghan, 2019). These results suggest that PlAMV tries to counteract plant defense by altering Ca²⁺ signaling.

GO enrichment analysis shows that ROS-related biological processes, such as hydrogen peroxide response and oxidative stress pathways, are strongly represented, while *RBOHD* is repressed. This suggests that redox balance is actively adjusted during infection. Since Ca²⁺ influx is linked to the activation of NADPH oxidases (RBOHs), which are the main source of apoplastic ROS (Kadota *et al*., 2015; Gilroy *et al*., 2016), the coordinated changes in Ca²⁺ and ROS genes point to a functional connection between these pathways in the plant’s response to PlAMV. GO enrichment of cellular components among downregulated genes shows that membrane-associated compartments are especially affected. Since many Ca²⁺ channels and ROS-producing enzymes are found in membranes, these changes in gene expression suggest that changes to membranes are closely tied to how signaling is reorganized during infection.

Overall, PlAMV infection leads to coordinated changes in membrane-associated Ca²⁺ and ROS signaling networks. This combined transcriptional response likely shows both the activation of the plant’s defense pathways and changes in host cell processes caused by the virus, highlighting the key role of the Ca²⁺–ROS axis in plant–virus interactions.

### Ca^2+^ signaling as an early antiviral response

For the first time, we employed genetically encoded biosensors to visualize host cellular signaling dynamics during viral propagation in living plants. Using a pioneering dual live-cell imaging method, we observed that Ca^2+^ signals originated at sites of virus-infected cells and propagated to neighboring cells before detectable accumulation of PlAMV–GFP in those cells. This temporal pattern strongly suggests that Ca^2+^ signaling is not simply a downstream consequence of viral replication but may also represent a host signaling event that modulates viral propagation.

Ca^2+^ influx is among the earliest host responses to biotic stress, acting upstream of multiple immune pathways (Thor *et al*., 2020; Negi *et al*., 2023). Our data mirror recent reports showing rapid, long-distance Ca²⁺ waves triggered by wounding or pathogen elicitors (Toyota *et al*., 2018; Wang *et al*., 2023). Such waves regulate reactive oxygen species (ROS) production, callose deposition, and transcriptional reprogramming (Then *et al*., 2021; Zvereva *et al*., 2024), suggesting that a similar mechanism may be engaged during PlAMV infection. The fact that Ca^2+^ elevation occurred specifically in PlAMV-infected cells but not in free-GFP–expressing controls underscores a virus-dependent trigger rather than a nonspecific fluorescence artifact (Figure 2 and Figure S5).

Our observation that Ca^2+^ signals precede detectable viral spread raises the possibility that PlAMV exploits or triggers Ca^2+^-mediated modulation of PD. PD are known to be regulated by Ca²⁺-dependent callose synthesis and degradation (German *et al*., 2023; Bayer & Benitez-Alfonso, 2024; Pérez-Sancho *et al*., 2025). Several plant viruses recruit host factors that alter plasmodesmata gating to enhance intercellular movement (Burch-Smith & Zambryski, 2016; Reagan & Burch-Smith, 2020). If PlAMV-induced Ca^2+^ waves open or prime plasmodesmata in neighboring cells, this could facilitate cell-to-cell movement of the virus. Conversely, Ca^2+^ spikes could represent a host defense mechanism attempting to close plasmodesmata, which the virus then needs to overcome. Distinguishing between these scenarios will require manipulating Ca^2+^ levels genetically or pharmacologically during viral infection. Consistent with the second hypothesis, mutants deficient in GLR3.3/3.6, and CNGC19 exhibited significantly larger infection foci compared to wild-type Col-0, suggesting that Ca^2+^ influx mediates viral propagation restriction. By contrast, mutants in the OSCA1 calcium channels displayed wild-type phenotypes. These results identify a specific subset of Ca^2+^-related components as negative regulators of PlAMV propagation. The combined transcriptional and imaging data thus highlight Ca^2+^ signaling as a rapid and spatially coordinated antiviral mechanism in Arabidopsis.

### Reactive oxygen species dynamics

PlAMV infection also modulates reactive oxygen species (ROS) homeostasis. Plasma membrane-localized ROS, visualized using the HyPer7-kRas biosensor, accumulated predominantly in cells distal to infection foci, indicating a spatially organized oxidative response. A similar pattern was observed using the cytosolic HyPer7 reporter, confirming that elevated ROS levels were not confined to the membrane compartment but reflected broader cellular redox changes.

Analysis of Arabidopsis mutants altered in ROS-producing NADPH oxidases RBOHD, RBOHF and RBOHD activator DGK5, revealed their critical role in limiting PlAMV spread. Both *rbohD* and *rbohF* mutants displayed significantly expanded infection foci, indicating that RBOH-mediated ROS production is essential for antiviral defense (Nadarajah, 2020; Kong *et al*., 2024). By contrast, the mutant in the LRR ROS sensor HPCA1 displayed wild-type infection foci, indicating that it does not contribute to the oxidative response restricting PlAMV propagation.

### Interplay between Ca^2+^ and ROS signaling during PlAMV infection

Ca^2+^ signaling frequently operates in tandem with ROS production and CPK activation at the plasma membrane (Kadota *et al*., 2014; Kadota *et al*., 2015). Our data showing altered ROS dynamics in *moca1*, *rbohD*, and *cpk3-2* mutants (Figures S5) support a model in which Ca²⁺ influx and NADPH oxidase activity are coordinated during early infection. Because CPK3 and RBOHD are classic Ca²⁺-responsive effectors (Dubiella *et al*., 2013), the early Ca^2+^ wave we detected may activate CPK3 through Ca²⁺ binding to its EF-hand motifs, which in turn phosphorylates and activates RBOHD. This cascade likely results in spatially patterned ROS accumulation and redox signaling that influence the dynamics of viral spread. The *cpk3/rbohD* double mutant has a larger PlAMV infection focus area than either single mutant, suggesting a synergistic interaction between CPK3 and RBOHD. This suggests that both proteins work in the same pathway or signaling module to help limit viral spread. Thus, Ca^2+^ and ROS signaling can function cooperatively to ensure robust antiviral response.

Although the *rbohD/moca1* double mutant displays the same phenotype as *rbohD* during PlAMV infection, this epistatic relationship does not necessarily imply that MOCA1 acts upstream of RBOHD in a single pathway. An alternative interpretation is that MOCA1-dependent lipid composition defines a permissive environment for viral propagation that is dominant over RBOHD-mediated restriction, such that disruption of lipid organization masks the contribution of ROS-based defenses. In this scenario, MOCA1/IPUT1 and RBOHD operate in parallel pathways, both influencing PD function but through mechanistically distinct processes.

The dominance of *moca1* in the *cpk3-2/moca1* double mutant further indicates that MOCA1-dependent PM sphingolipid represents a downstream node where Ca^2+^ and ROS signaling converge. The similarity in infection phenotypes between CPK3 and RBOHD mutants suggests a functional link between Ca^2+^ signaling and localized ROS production in antiviral defense. CPK3 may act upstream of or coordinate with RBOHD to fine-tune ROS-mediated viral restriction. *moca1* mutants, defective in the Salt Overly Sensitive (SOS) pathway and hypersensitive to salt stress (Jiang *et al*., 2019b), failed to spatially constrain ROS accumulation during virus propagation, implicating membrane sphingolipid in the regulation of ROS production/localization during virus propagation (Figure S7A and D). Collectively, these results establish a mechanistic link between Ca²⁺ signaling, RBOH activation, and ROS-mediated defense that shapes the spatial and temporal dynamics of viral propagation.

Taken together, our findings support a model in which RBOHD-mediated ROS production functions either in coordination with MOCA1 or through an independent pathway, indicating that ROS signaling and plasma membrane sphingolipid homeostasis are interconnected in antiviral defense, while CPK3 contributes an independent Ca²⁺-mediated signaling pathway. The distinct and sometimes opposing effects of these components on local viral spread emphasize that ROS and Ca^2+^ signaling operate in a spatially coordinated but mechanistically diverse manner to maintain antiviral defense homeostasis.

## Perspectives and Future Directions

Our work identifies GLR3, CNGC and RBOHD as critical components of the host response to PlAMV infection. Despite these advances, key questions remain. How does PlAMV modulate Ca^2+^ channels and ROS accumulation at the plasma membrane? Which Ca^2+^ channels are responsible for the Ca^2+^ wave observed in the cells surrounding the infection foci?

Future studies could address these questions using complementary approaches. Super-resolution imaging and single-particle tracking could resolve Ca^2+^ and ROS nanodomain dynamics at subcellular scales. Moreover, pharmacological inhibition or genetic disruption of Ca²⁺ channels, and pumps will be crucial to test whether blocking the early Ca^2+^ wave impairs viral spread. Finally, high-resolution analysis of PD ultrastructure in the presence or absence of Ca²⁺ flux will clarify how Ca^2+^ signaling interconnects with intercellular transport.

## Conclusions

Overall, our findings support a model in which PlAMV infection triggers a rapid, spatially organized Ca^2+^ wave that precedes viral spread. This early signal seems to coordinate ROS production and shape the local environment, possibly preparing nearby cells for virus propagation. Ca^2+^ and ROS signaling work together to limit viral propagation between cells. These results highlight the importance of early membrane signaling networks in plant antiviral defense and offer a basis for improving viral resistance in crops through the targeted gene editing of Ca^2+^ and ROS pathways.

## Data availability

The submission of RNAseq data to GEO is in progress (Edgard R. *et al*., 2002, http://www.ncbi.nlm.nih.gov/geo/). All steps of the experiment, from growth conditions to bioinformatic analyses, were detailed in CATdb database (Gagnot *et al., 2007*, http://tools.ips2.u-psud.fr/CATdb/) under project name 2022_10_QS_PhosphoRem_B1.

## Acknowledgements

We thank Thierry Mauduit (HPE Greenhouse, INRAe) for plant culture. This work was supported by the European Union’s Horizon 2020 research and innovation program under a Marie Skłodowska-Curie grant (grant no. 101104279 to JA) and the French National Research Agency (grant no. ANR-24-CE20-7620 and 11–INBS–0010 to SM, VG, MB). The IPS2 and the POPS platform benefit from the support of Saclay Plant Sciences-SPS (ANR-17-EUR-0007). We thank the Bordeaux Imaging Center, part of the National Infrastructure France-BioImaging supported by the French National Research Agency (ANR-10-INBS-04). We are grateful to Ron Mittler for the *glr3.3/glr3.6* mutant lines, to Cyril Zipfel for the *osca1;1/1;2/1;3, bik1, bak1-5/bkk1-1*, and *serk1* lines, and to Zhen-Ming Pei for the *hpca1* and *moca1* lines. This study received financial support from the French government in the framework of the IdEX Bordeaux University “Investments for the Future” program / GPR Bordeaux Plant Sciences

## Notes

### Competing Interest Statement

The authors have declared no competing interest.

