## Supplementary figures and images for "The role of reactive oxygen species and calcium signaling in antiviral defense in Arabidopsis"

### Figure S1

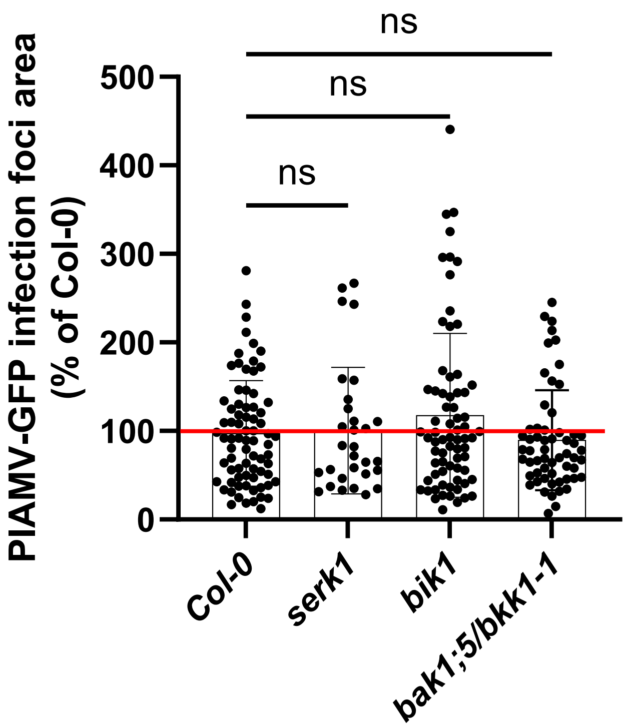

### Figure S2

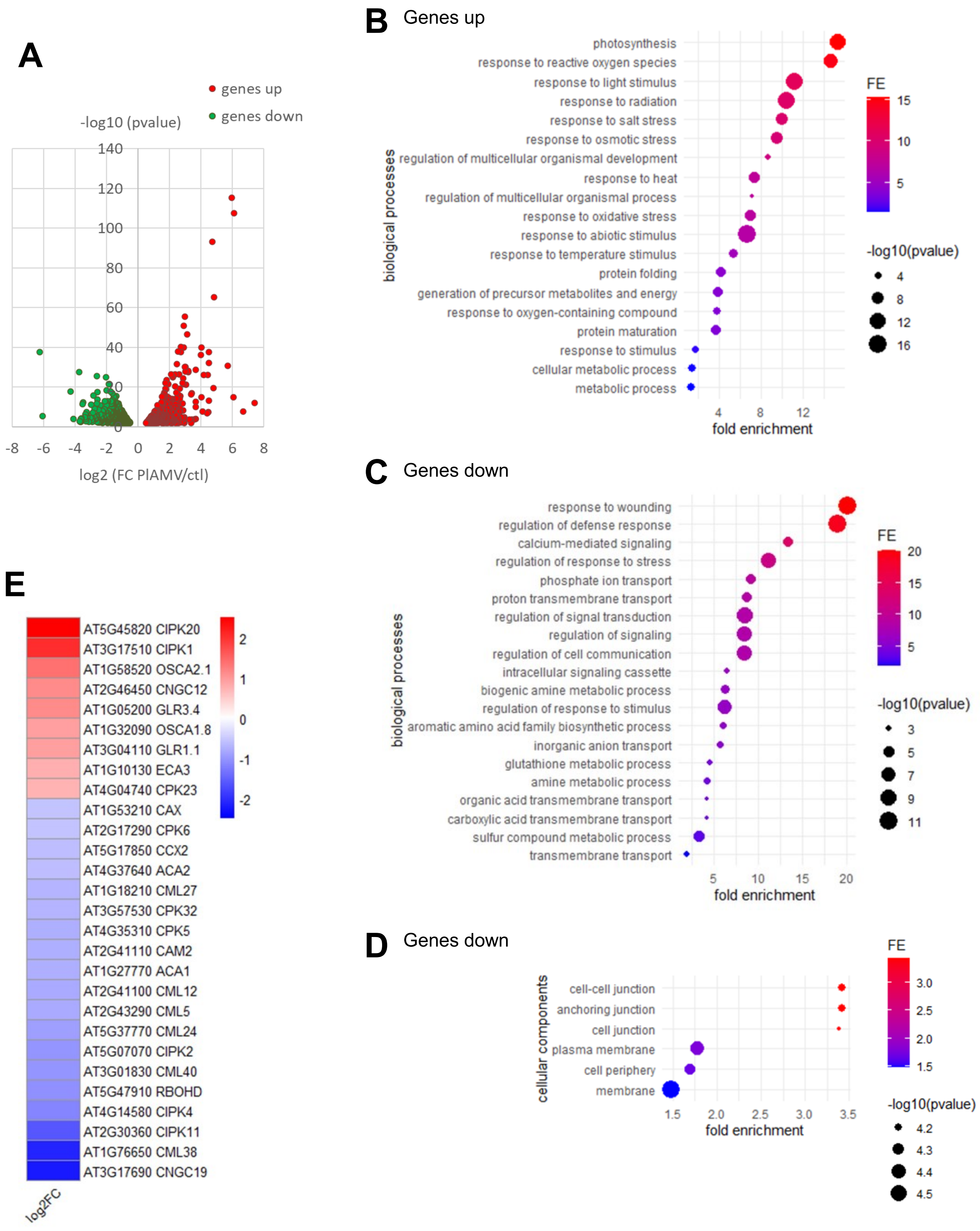

### Figure S3

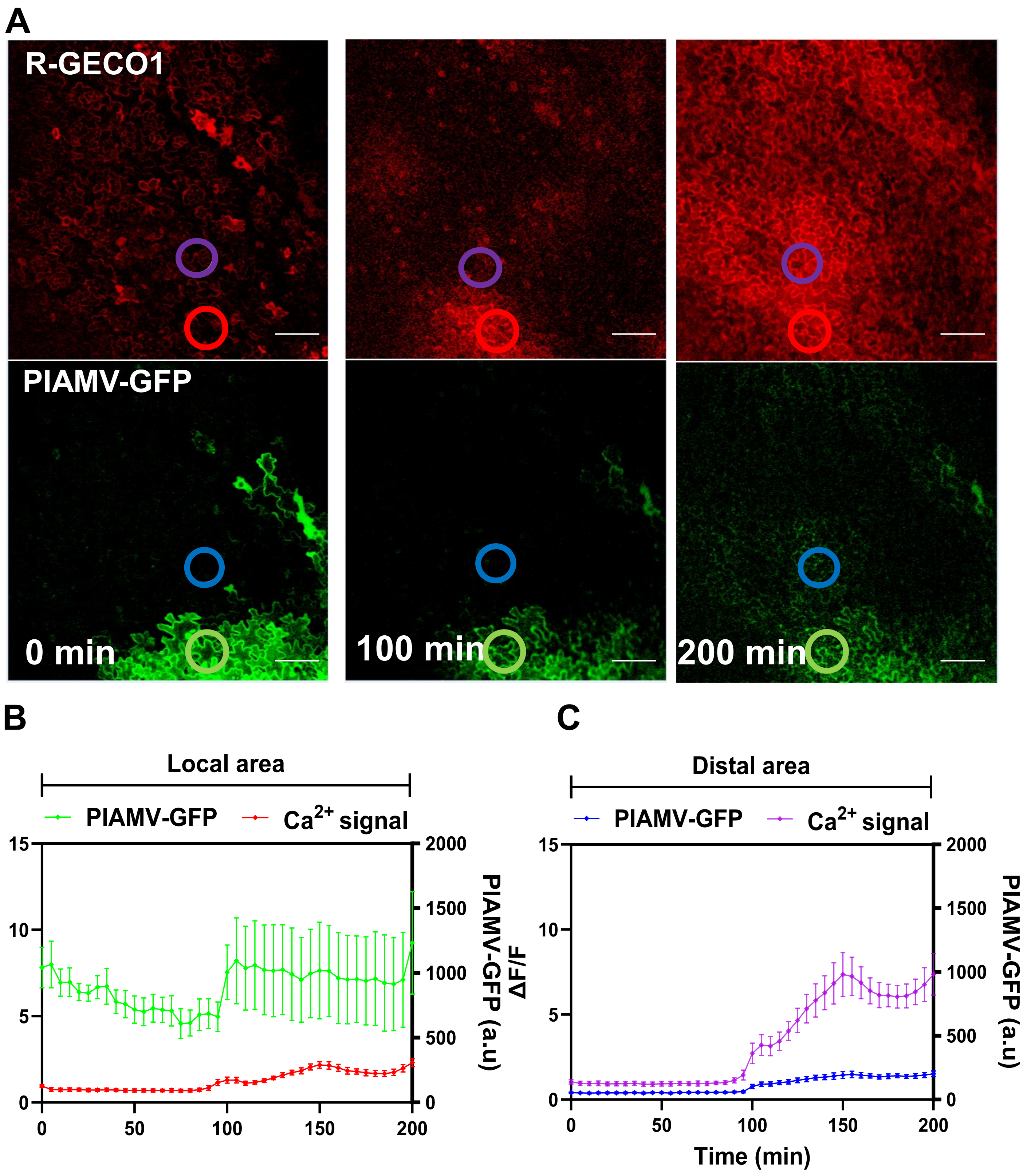

### Figure S4

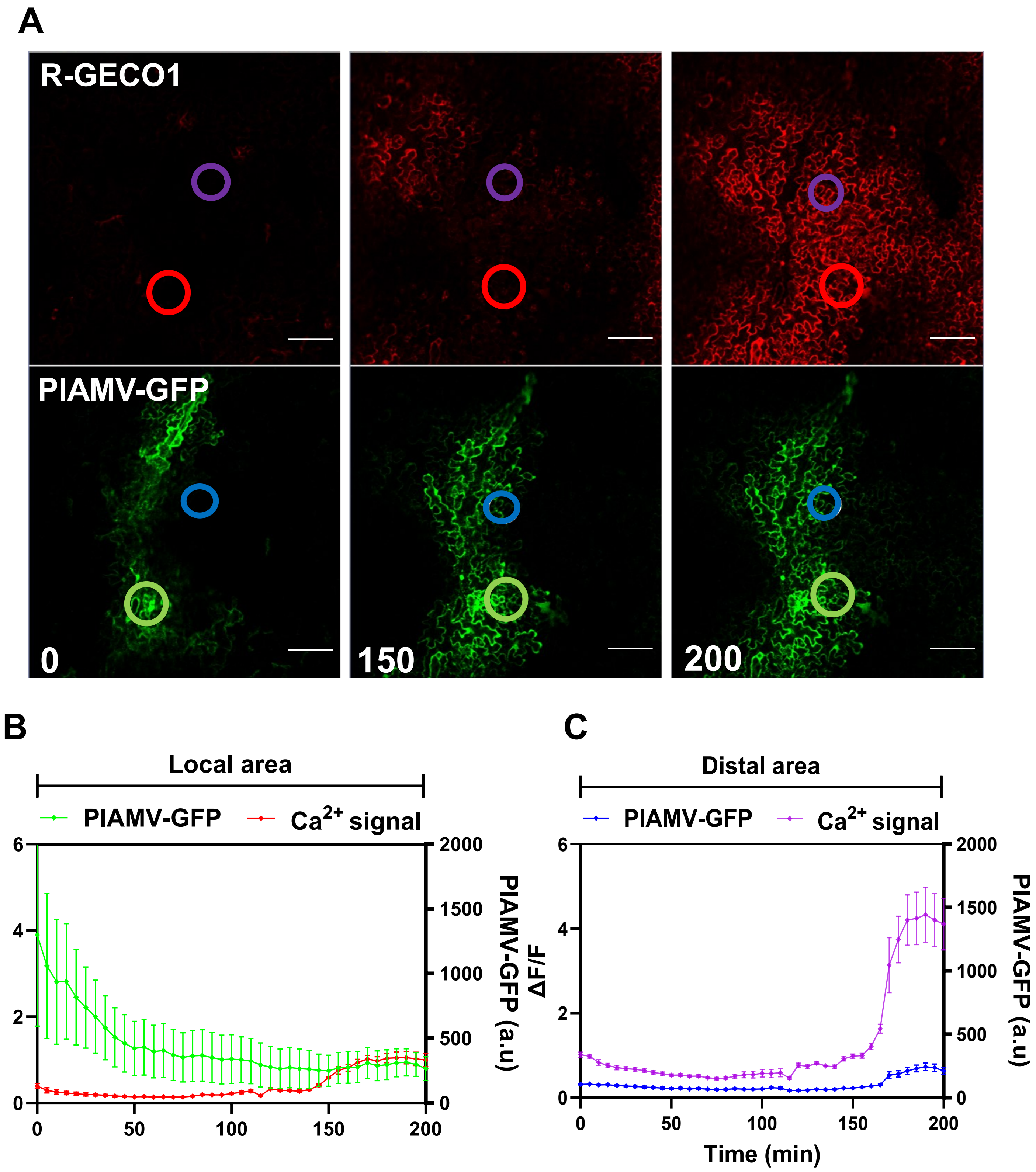

### Figure S5

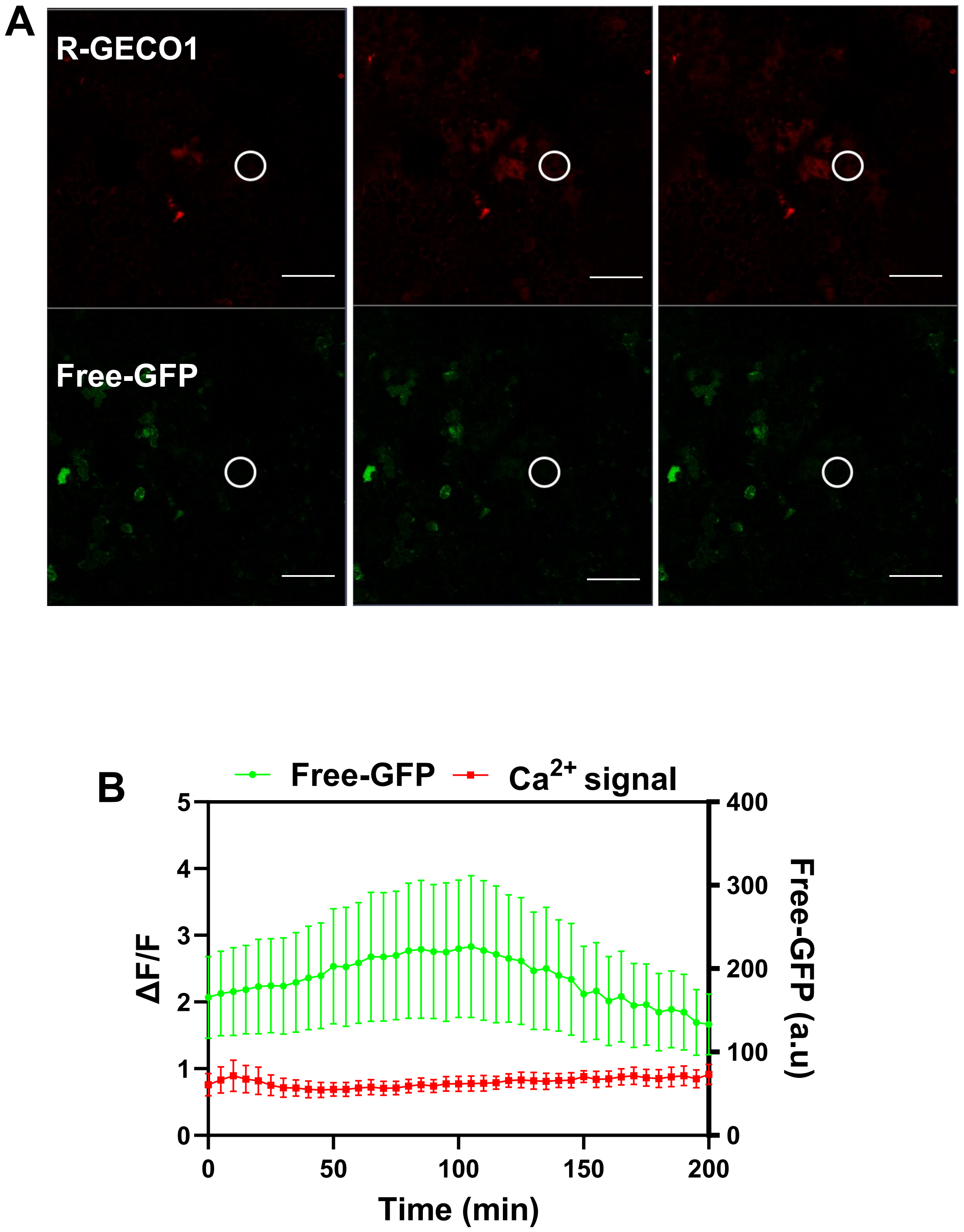

### Figure S6

Fig S6

A

***HyPer7-kRas***

**405** **488**

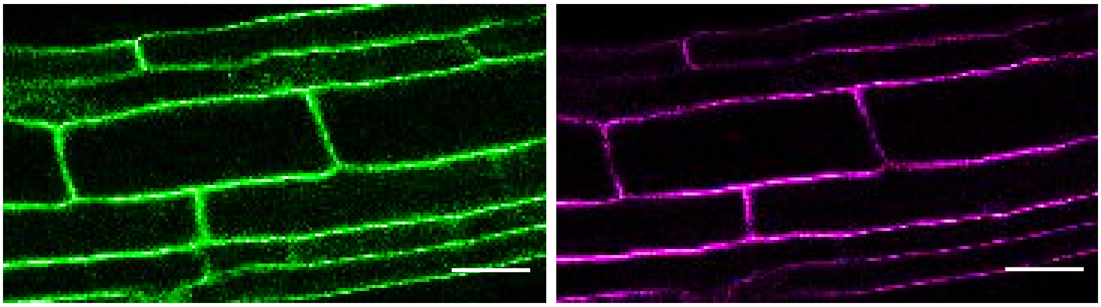

B

***HyPer7-Cyto***

**405** **488**

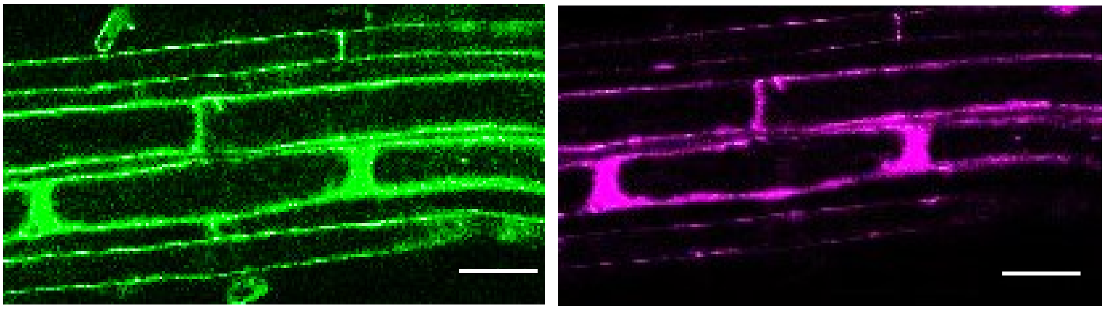

### Figure S7

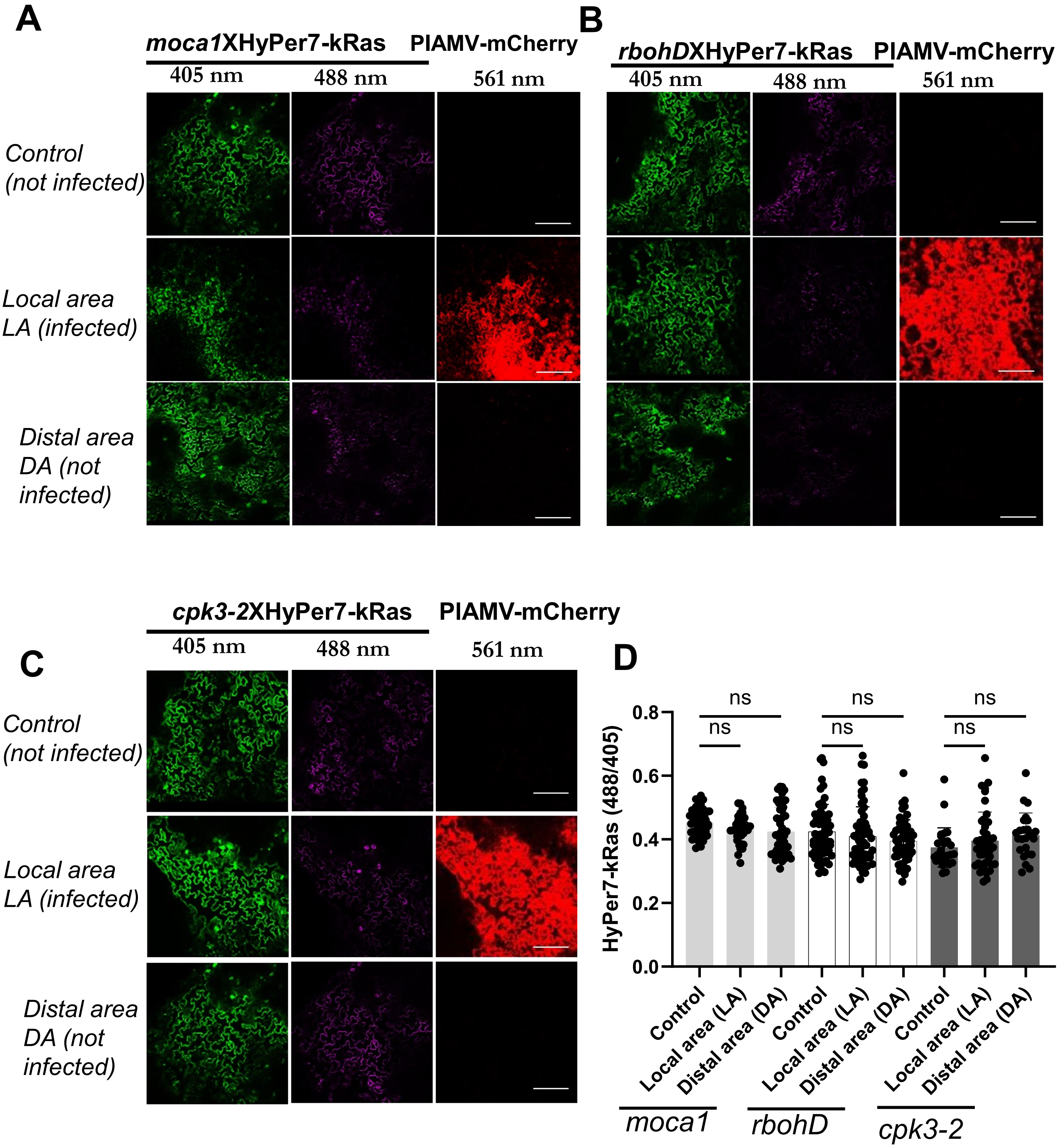
